# Variation in spatial dependencies across the cortical mantle discriminates the functional behaviour of primary and association cortex

**DOI:** 10.1101/2023.01.13.523934

**Authors:** Robert Leech, Reinder Vos De Wael, Frantisek Vasa, Ting Xu, R. Austin Benn, Robert Scholz, Rodrigo M. Braga, Michael Milham, Jessica Royer, Boris Bernhardt, Emily Jones, Elizabeth Jefferies, Daniel Margulies, Jonathan Smallwood

## Abstract

Recent theories of cortical organisation maintain that important features of brain function emerge through the spatial arrangement of regions of cortex. For example, areas of association cortex are located in regions of cortex furthest from sensory and motor cortex. Association cortex is also ‘interdigitated’ since adjacent regions can have relatively different patterns of functional connectivity. It is assumed that topographic properties such as distance between cortical regions constrain their functions. For example, large distances between association and sensory and motor systems may enable these areas of cortex to maintain differentiable neural patterns, while an interdigitated organisation may enable association cortex to contain many functional systems in a relatively compact space. We currently lack a formal understanding of how spatial organisation impacts brain function, limiting the ability to leverage cortical topography to facilitate better interpretations of a regions function. Here we use variograms, a quantification of spatial autocorrelation, to develop a cortex-wide profile of how functional similarity changes as a function of the distance between regions. We establish that function changes gradually within sensory and motor cortex as the distance between regions increases, while in association cortex function changes rapidly over shorter distances. Subsequent analysis suggests these differential classes of spatial dependency are related to variation in intracortical myelin between sensory motor and association cortex. Our study suggests primary and association cortex are differentiated by the degree to which function varies over space, emphasising the need to formally account for spatial properties when estimating a system’s contribution to cognition and behaviour.

**Significance statement:** The spatial arrangements of regions in the human brain are hypothesised to underpin important features of a brain regions function. Currently, however, we lack a formal understanding of how topography shapes brain function, limiting our ability to leverage topographical perspectives to inform better theories of brain function. Here we use a formal mathematical approach to establish that in regions of association cortex function varies across the cortex more rapidly than in sensory and motor cortex, a phenomenon linked to levels of intracortical myelin. This result highlights how topographical features distinguish between cortical regions with different functional profiles and provides a formal account of how spatial differences support different features of brain function.

## Introduction

One of the most important discoveries in human neuroscience is that brain topography plays an important role in determining how a region contributes to cognition and behaviour (1). These topographic features can shape a region’s function in many ways including: (i) through the influence of neighbouring neural systems that make up the local environment within which a specific region is embedded (2), (ii) the physical location of the network on the cortical mantle with respect to core cortical landmarks (3), (iii) and more abstract topographical features such as the degree to which functional activity within a network is spatially distributed across the cortical mantle (2, 4), or, instead is limited to adjacent regions, often within a single cortical lobe (5, 6).

Contemporary evidence suggests that local topographical properties influence a region’s function in a complicated, interdependent manner. For example, neural systems that are directly concerned with sensation and movement, such as the visual or motor cortex, are spatially distant from each other, yet both of these systems tend to be relatively spatially contiguous, and both contain topographic features resembling maps, either of the external environment or how the organism engages with the outside world (7–10). Other systems, such as the default mode or frontoparietal networks, are located in regions of association cortex, are spatially adjacent to one another, and both are spatially distributed across cortex; yet functionally these systems appear to serve different, often opposing roles in human cognition (11). Topography is also important for understanding macroscale brain function, because systems that tend to be more spatially discontinuous (e.g., the default mode network) tend to be more distant from sensory and motor systems where spatial discontinuity is an exception rather than the norm (e.g. sensorimotor or visual cortex) (3). In contemporary neuroscience, macroscale topographical features provide a useful heuristic for understanding the involvement of frontoparietal and default mode networks in cognition. These networks are hypothesised to be at the transmodal apex regions of a broad sensory-fugal hierarchy, allowing oversight across broad areas of cortex (12). In contrast, mesoscale features of topography, such as the retinotopic maps located within sensory cortex, are thought to explain aspects of how the visual system represents and extracts features of the environment from retinal input.

Topography at both macro and mesoscale is, therefore, a key principle of brain organization and is crucial for understanding brain function both within specific systems and across the cortex as a whole. Our study set out to formally examine how this perspective can be leveraged to go beyond heuristic accounts of the topographic influence of a region’s contribution to cognition and behaviour. The distance between regions, calculated as the *geodesic distance* between two vertices, provides one metric to understand the influence of topography on neural function can be approached. This measure has been used to show that systems like the default mode and frontal parietal cortex are both distant from the sensory input and motor output systems (13). However, a given location on the cortical mantle may be influenced by topography in a number of ways, such as through features of the local neighbourhood in which the region is situated, or, whether the system is part of a distributed or localised network. Accordingly, it is important to understand whether the way that distance between two regions influences neural function varies across the cortex. Understanding if there are regional differences in the way distance impacts functional connectivity (second-order non-stationarity), enables us to leverage a topographic perspective to better understand neural function.

In order to establish how distance between regions influences their similarity in function, we calculated for each cortical vertex surface how the similarity of its activity changes with all other vertices as a function of the distance between them. This is a simplified version of the empirical variogram (14), as illustrated schematically in the upper panel of Figure 1. Spatial variograms are expected to show that similarity in function declines with distance until it reaches an asymptote, the distance after which there is no longer a spatial dependency between vertices. The empirical variogram can be summarised by fitting an exponential function which in turn can be described by two values capturing how similarity changes with distance for each vertex: the effective range and the sill. The sill is the height (i.e., degree of dissimilarity between two regions) and the range is asymptote (i.e., the spatial distance between the two regions). Differences in the shape of the spatial variogram across regions can be used to quantify the different ways topography influences function in different cortical locations. For example, in regions where function is more influenced by the local neighbourhood, the spatial variogram shows a relatively shallow decline in similarity with distance. In contrast, in regions where function is relatively distinct from the local environment, the variogram should increase more rapidly with the distance.

**Figure 1.**
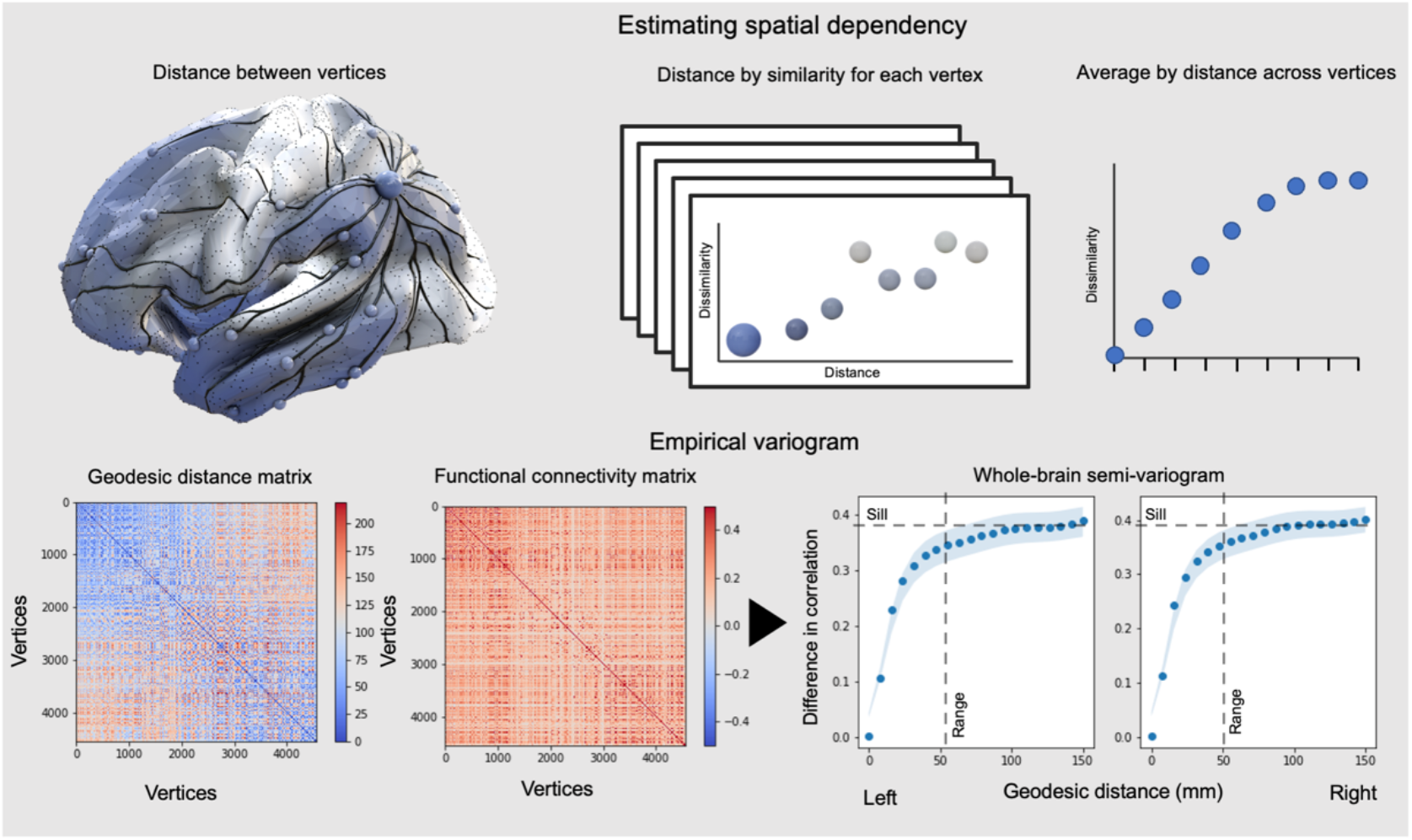
Top Panel. Schematic illustration of how spatial variograms can be used to characterise the rate of change of functional connectivity as distance increases between brain regions. **Bottom Left**. Whole-brain variograms of functional connectivity can be calculated by comparing how the distance along the cortical surface is related to the average similarity in brain activity between regions. **Bottom Right**. Whole brain variograms are shown for the left and right cortices and can be seen to be broadly similar. The thick lines/dots are the mean across participants, and the filled area is the standard error of the mean. The dashed lines are the estimated location of the sill (asymptotic difference in correlation between vertices) and range (distance in mm between vertices at which the asymptote is reached).

## Results

We first quantified the spatial dependency between functional connectivity and distance by calculating whole-brain variograms assessing how functional connectivity affinity matrices (Pearson’s correlation) vary with distance along the cortical surface for each hemisphere (Figure 1, lower panel). We used resting state fMRI data from 51 participants from the Human Connectome Project for whom there are two sessions separated by approximately six months. This analytic choice allows us to calculate the reliability of these metrics across time within an individual. Averaging these vertex-wise variograms across the whole cortex, the global variogram, reveals an initially steep rise (rapidly increasing dissimilarity with distance), with an inflection point at around 30-40 mm (Figure 1). This is followed by a continuous increase up to the limit of measurable distance (all vertices included distances up to 150 mm; however, measurements above this value are not present in all vertices and so become more unstable and unduly influenced by a subset of regions). In other words, at the level of resolution of the present data, vertices within <40 mm of each other show similar temporal profiles of activity. The variogram for the left and right hemispheres show a similar pattern (see left hand panel). The landscape of these variograms can be formally understood by comparing the observed rate of change in function with distance with different mathematical growth functions (e.g., sinusoidal, exponential, or gaussian). It can be seen in Figure 1 that the whole brain variogram of the human is most similar to an exponential relationship.

For the purposes of our analyses, we can extract the two parameters used to fit the theoretical function to the empirical variograms: (i) the sill is where the height of the variogram reaches 95% of its asymptote; and (ii) The distance where the sill occurs defines the range. These are both displayed in the top panel of Figure 2. Importantly comparing the variogram calculated for each of the participants on two resting state sessions separated by 6 months shows a high degree of correspondence both in terms of the sill (the average difference in correlation between vertices) and the distance (i.e., rho > 0.73; Figure 2 top panel).

**Figure 2.**
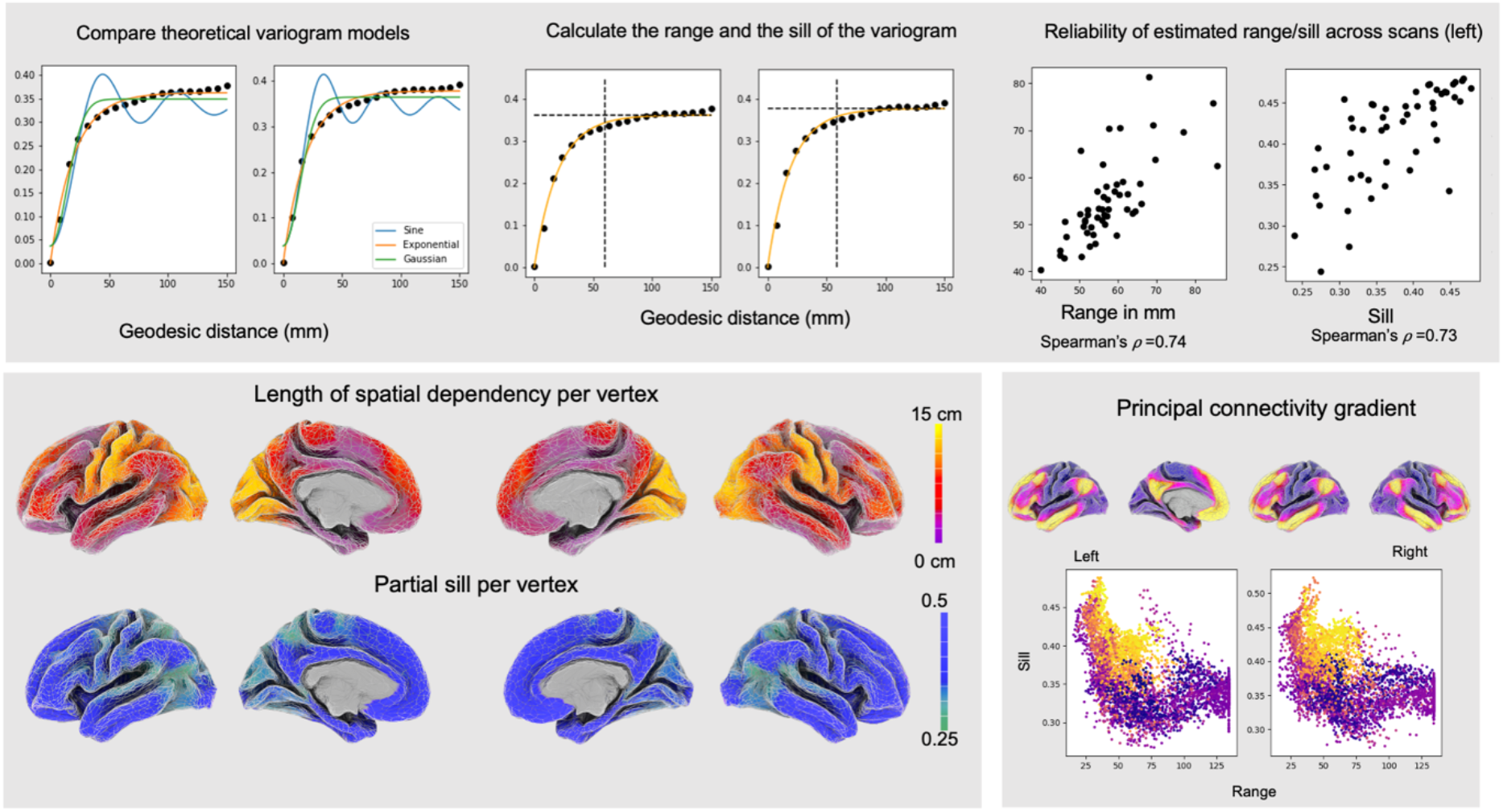
Distribution of the sill and effective distance of variograms across the cortex. **Top Panel**. Variograms can be formally described through comparison of the observed rate of change between similarity in brain activity and distance with different mathematical growth functions. We observe that the whole-brain variogram has most similarity to an exponential function. **Top Middle**. Variograms can be described in two numbers, the partial sill (the height of the curve at 95% of its asymptote) and the effective range (the distance of the sill). **Top Right**. Both the partial sill and the effective range of the whole brain variogram show reasonable similarity when measured within the same individual at two different timepoints (> 0.73). **Bottom left**. The distribution of the partial sill (height of the variogram at 95% of its asymptote) and effective distance (the distance of the sill) across the vertices of the human cortex. It can be seen that the sill ranges from .25 and .5 across the cortex and that in some regions effective distance can be as long as 15 cm. **Bottom right**. The relationship between the distribution of the principal gradient of intrinsic connectivity and variograms at each vertex (as described by each vertex’s partial sill and effective distance). It can be seen that unimodal regions tend to have variograms with low sills and long effective distances, while transmodal regions have variograms with higher sills and shorter effective distances.

The whole-brain variograms establish that in humans, distance leads to an increase in dissimilarity in neural function that is asymptotic exponential in nature and that these measurements are broadly consistent within an individual over time. This aligns with descriptions of local spatial similarity previously reported in humans and non-human primates e.g., (15). By computing variograms, we are able to go beyond a single description of spatial dependency across the brain, and capture regional differences in spatial dependencies (i.e., second-order non-stationarity). To understand whether there are systematic differences in how distance leads to changes in neural function across different brain regions, we calculated separate variograms for each vertex across the cortex. The middle panel in Figure 2 summarises how the two metrics (sill and effective distance) vary across the cortex. It can be seen that sill (reflecting the spatial dissimilarity in functional connectivity across the cortex) ranges between 0.25 and 0.5, and that in some regions the dissimilarity continues to increase to the maximum range of our measurements (150 mm).

To further characterize this heterogeneity, we examined how the distribution of the sill and the effective range varies with the principal gradient of change in functional connectivity (3). This gradient can be derived by application of dimensionality reduction techniques to functional connectivity data (3), and recapitulates foundational models of the sensory-transmodal cortical hierarchy (1). The lower panel of Figure 2 shows that regions closer to the transmodal end of the principal gradient tend to be regions where the variograms have a relatively high sill and short effective distance (i.e., regions where dissimilarity shows a relatively rapid increase). In contrast, regions closer to the unimodal end of the principal gradient have a relatively lower sill and a longer effective distance (i.e., regions that show a slower rate of decline in function with increasing distance). This analysis provides preliminary support that two broad types of cortex (primary and association cortex) can be discriminated based on how activity varies with distance.

Next, we calculated spatial dependency for each of the canonical resting state networks to understand whether this is true of different large-scale networks within both association and primary cortex. Figure 3 (upper panel) shows the average empirical variogram for each network while the lower panel shows the average sill and effective distance of each network. Regions making up the limbic network (Cream) have the highest sill and the shortest effective distance, a pattern that is also seen in the transmodal networks (Default mode, Red; Fronto-parietal network, Orange) but to a lesser degree. Regions that make up unimodal cortex (Visual network, Purple; Motor cortex, Blue) show the reverse profile with variograms with small sills and relatively long effective distance. Finally, the two attention networks (Dorsal and Ventral) show intermediate profiles both having moderate sills and effective ranges. These two systems are distinguished from each other because the Dorsal attention network has a longer effective distance and a short sill, and so is more similar to the unimodal systems, whereas the Ventral attention shows the opposite profile.

**Figure 3.**
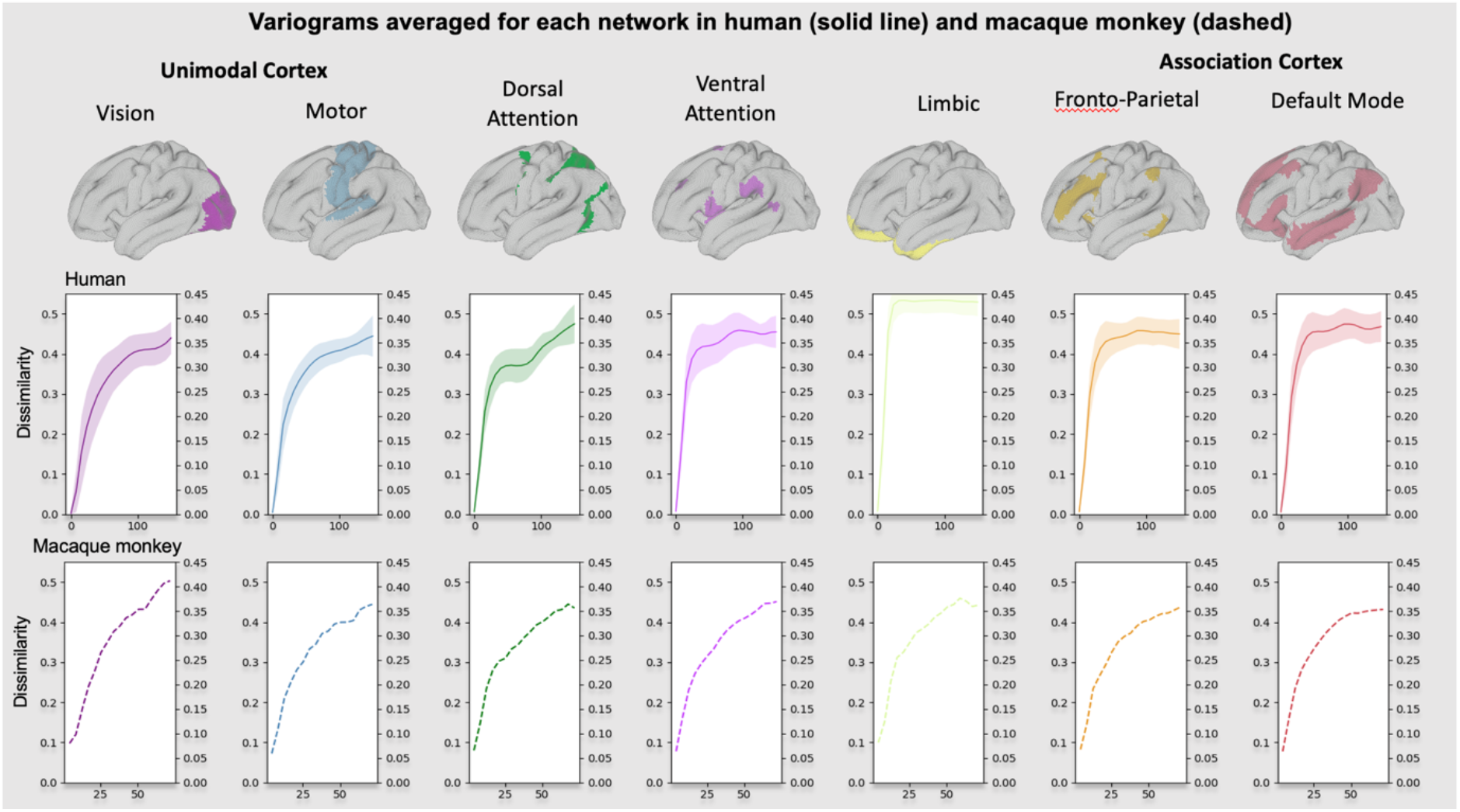
Variograms calculated for each canonical resting state network (Yeo, Krienen et al., 2011) in humans and in homolog networks in macaques. The middle panel shows the mean variogram (FC dissimilarity by distance along the cortex) calculated across all vertices for each Yeo network in the human Human Connectome Project data; the filled areas are the standard errors of the mean across vertices. The bottom panel shows a similar analysis with fMRI data averaged from 14 awake Macaque monkey as a comparison.

We repeated this analysis in a sample of macaques to determine whether these properties are unique to humans (using homolog networks, see Methods for details). This analysis identified that the network profile of each species is broadly similar. For example, in both species the Limbic network has the highest sills and the shortest effective distances, and the visual system provides the clearest example of the opposite profile (low sills and longer effective distance).

The variograms stratified by resting-state network suggest that there may be a small set of spatial dependency profiles that characterize a larger number of networks, and that these likely correspond to the difference between association and primary cortex. To provide an independent test of this idea, we performed hierarchical clustering on the binned data from the vertex-wise variograms, and display the results colored by different canonical networks. The top panel of Figure 4 presents the dendrogram produced by this analysis. Clustering vertices based on their variogram profiles gives rise to two groups, one predominantly encompassing the unimodal systems (primary sensorimotor networks as well as parts of the dorsal attention network) and the other corresponding to limbic and transmodal systems, as well as the ventral attention network. This analysis, therefore, highlights a broad dissociation of cortex into two classes based on their variograms: one class of regions where the variograms have low sills and long connectivity and a second class of regions with higher sills and shorter effective distances. We also assessed how consistent these results were for individuals’ variograms across different scans (Figure 4C), to ensure the cluster structure was not a consequence of group averaging and generalises to out-of-sample data. Comparing each individual participant’s empirical variograms across scans showed within-cluster correlations (cluster variograms from scan 1 correlated with cluster variograms from scan 2 substantially higher than across clusters).

**Figure 4.**
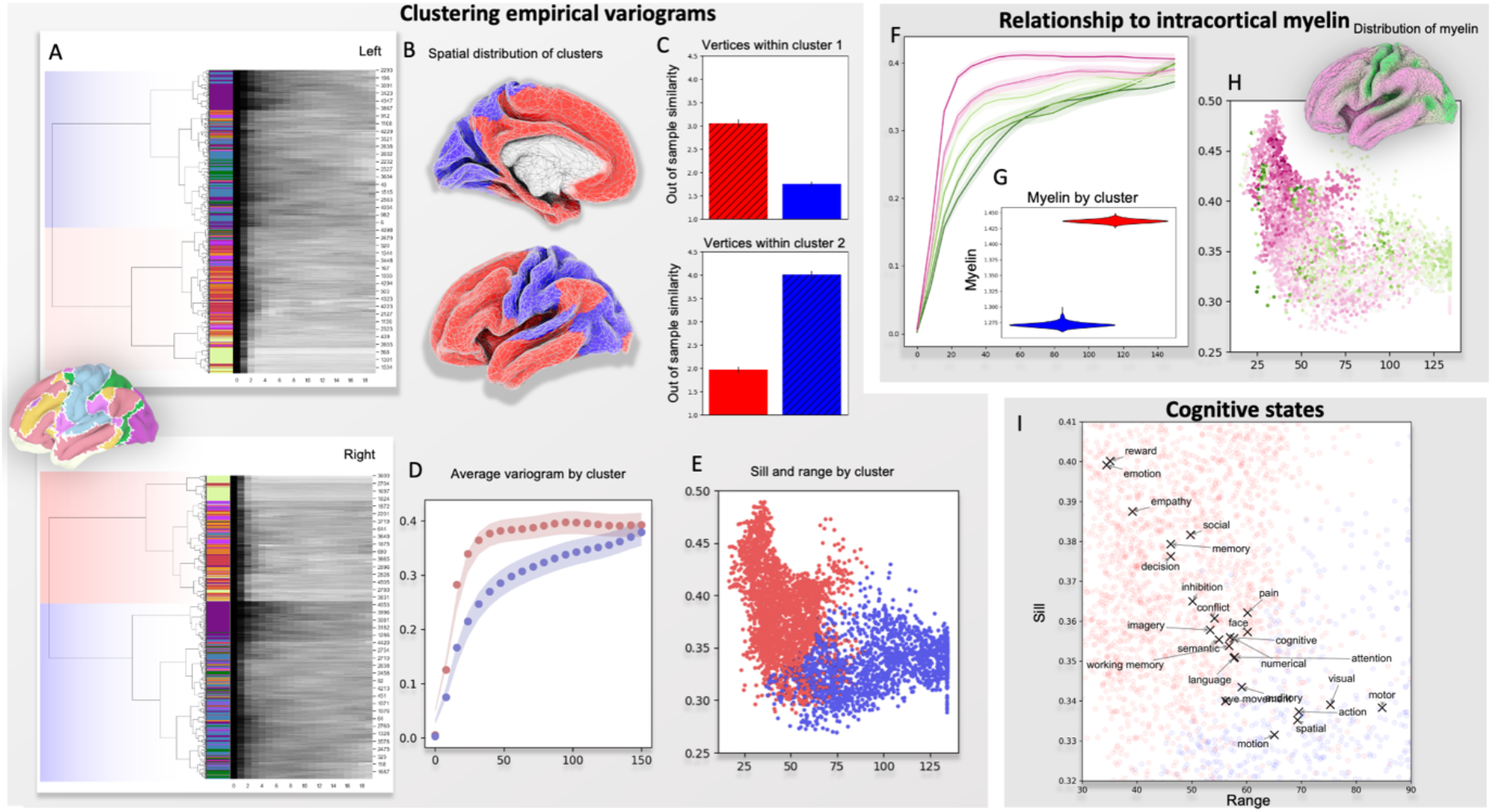
Left. A: Clustering vertices based on empirical variograms. The dendograms, are colored by the Yeo network that each vertex belongs to, displaying the tree structure of the similarity between variograms. B: The dendogram was used to cluster the data into two clusters (colored red and blue) for the left and right hemispheres. The order of the clustering was arbitrary across hemispheres and has been colored based on approximate similarity between the left and right hemispheres. Broadly, transmodal regions were clustered together in a separate cluster (red) to unimodal sensorimotor regions (blue). C: Correlation of empirical variograms across vertices are consistent within each cluster within individuals and across different MR sessions. D: Average empirical variograms for each of the clusters within individuals reveals that one cluster exhibits more dramatic change in functional similarity with distance. E: The range and sill for each vertex, colored by the cluster label for the left and right hemispheres. **Top, right**, F: The empirical variograms between functional connectivity and distance split into deciles based on vertices’ myelin value (pink-greener colors correspond to higher- myelin content). G: Individual average estimated intracortical myelin for the two clusters. I: The estimated range and sill for each vertex, colored by estimated myelin. The inset brain is the average distribution of estimated cortical myelin (from the HCP group average dataset). **Bottom, right**, the ranges and sills calculated across vertices activated by different cognitive processes (taken from a large automatic meta-analysis); These are overlayed on vertices colored by their cluster membership from E.

Given the different profiles of spatial dependencies observed across different regions of cortex, we investigated whether the distribution of activity for different cognitive states also reflects this. To this end, we averaged vertex-wise estimates of the range and sill parameters for responsive vertices (defined as those with an estimated evoked BOLD response greater than threshold) in 24 topic maps generated by an automatic meta-analysis of functional MRI tasks. Figure 4 shows how brain regions related to different cognitive states differ in terms of their profile of spatial dependencies. In general, more externally focused tasks (e.g., labelled “visual” or “motor”) showed slower decrease in similarity with distance and a lower sill; whereas cognitive tasks associated with more abstract functions (such as “emotion”, “social”, “memory”), were associated with the opposite pattern with shorter ranges and higher sills. We subsequently clustered the tasks according to their sills/ranges to allow us to easily visualise the variability in the variograms associated with each task (the red/blue colors in Figure 4, panels A-E). This allowed us to create a composite task activation map for each cluster and plot the associated variograms showing the different spatial dependency profiles.

Our final analysis examined how microstructural features of different regions of the cortex correspond to the observed differences in spatial dependency profiles across cortex. Given its role in signal propagation, we examined whether myelination is linked to the shape of the variograms for different vertices. Figure 4H depicts the spatial distribution of estimated cortical myelin. We split vertices into deciles based on their levels of cortical myelination and plotted separate variograms for each decile. A clear separation emerges, with more highly myelinated vertices displaying, on average, longer distance spatial dependencies, and lower sills. This is made more explicit by plotting the range and the sill per vertex (Figure 4) colored by the level of myelination (warm colors indicating higher myelination).

## Discussion

Given emerging evidence of the importance of topography in the mammalian cortex (3, 12), our study set out to understand how the distance between regions relates to their functional similarity. In particular, we examined whether this profile of spatial dependence varies across different cortical regions (a phenomenon known as *second-order non-stationarity*). Our analysis first established whole brain variograms are reasonably consistent across hemispheres, individuals, and within individuals measured at different time points. When we examined these on a regional basis, we observed substantial differences that reflect known functional divisions of brain function. Notably, the observed differences in spatial dependence profile recapitulated the distinction between primary sensorimotor and transmodal association cortex. In primary sensorimotor cortices, including visual and somatosensory cortex, we found that increasing distance is associated with a gradual change in function. In contrast, in association cortex we found that function changed with distance at a much faster rate. These differences between unimodal and association cortex in humans were broadly similar to those seen in macaques suggesting that they are conserved across the primate nervous system. We found that these changes in how distance impacts functional variation are likely to be at least partly related to differences in microstructure, as we found differences between association and unimodal cortex similar to those seen when exploring variation in intracortical microstructure approximated by the ratio of T1w/T2w image intensity a known proxy for intracortical myeloarchitecture (16).

These results have implications for understanding how topographic differences influence cortical function. First, our data provides novel support for an organisation of unimodal cortex that supports the progressive elaboration of encoded stimulus features (17). Our analysis established that both sensorimotor cortex and visual cortex are situated within regions in which the changes in function over distance are some of the most gradual when the cortex is viewed as a whole. When contrasted with association cortex, this pattern is consistent with the view that sensory regions have a spatial organisation in which adjacent regions encode progressively complex features of the information extracted from sensory signals and that these compressed signals form the basis of signal processing for the next stage in the hierarchy e.g. (18). This pattern of progressive change is assumed to be important in regions of primary cortex, such as visual cortex, and is captured empirically by the variograms in these regions which show relatively small steady changes in functional properties as the distance between two regions increases.

Our study also provides insight into theoretical perspectives on how neural processing occurs in regions of association cortex. For example, contemporary work highlights that regions of association cortex can have relatively unique features both in terms of the functions they support, and in their observed neural properties (for a similar argument see (12)). For example, both the fronto-parietal and default mode networks are implicated in cognition in a relatively abstract manner, highlighted by their involvement in a wide range of tasks which despite being superficially different may draw on similar underlying cognitive operations. For example, situations which have superficially different features, such as the Stroop (19) or working memory (20), but show a common reliance on executive control, tend to activate the fronto-parietal network, as well as other task positive systems (21). Similarly, the default mode network is often observed as contributing to situations when information from memory may be important for organising cognition, such as during mental time travel (22), memory processes that rely on semantic (23) or episodic knowledge (24). Our analysis suggests that both of these large-scale systems are situated in regions of cortex where there are fairly rapid changes in functional similarity with increasing distance. These rapid changes in function over relatively short distances are likely to reflect the interdigitated nature of these systems (6, 25). These perspectives assume that a general property of associative cortex may be a topographic organisation in which relatively different functional systems terminate within close proximity of one another. This topographic system could form the basis of an architecture that is hypothesised to explain why both the fronto-parietal (26) and default mode networks (12) contribute to multiple different forms of behaviour in a relatively abstract manner. These more complex, interdigitated patterns of function are captured empirically by the variograms which show rapid functional changes as a function of distance in each of the large-scale networks in association cortex.

Finally, our study provides insights into the important observation that the default mode network, a brain system located at the maximal distance from primary landmarks like the calcarine sulcus, also has a functional profile which is one of the most unique in the mammalian nervous system (3). Our analysis suggests regions of cortex where the default mode network is located combine two unique topographic properties that together explain why the distance between these systems and the primary sensorimotor landmarks corresponds to the primary dimension of functional differentiation with the whole brain connectivity space (3). Our analysis suggests that the increasing distance from primary landmarks in sensory cortex, and regions of the DMN would first lead to increasing differences in functional similarity through the slow progressive changes in function with distance that emerge in primary cortex. In conjunction, with these gradual changes, our study suggests that the cortex where the DMN is where function changes most rapidly with increasing spatial distance. Thus, the observation that the distance between the DMN and sensory cortex corresponds to the greatest differentiation in function (i.e. the principle gradient of functional connectivity (3)) is inevitable because this distance combines (i) the progressive changes in function within primary sensorimotor cortex, and (ii) the complex interdigitated structure seen within the DMN (6). Based on our analysis of T1w/T2w images it is possible that microstructural differences, such as myelin content, may be an important feature in distinguishing these types of cortex, an important question for future research to explore with more detailed anatomical techniques (e.g., (27) than those used in the current investigation.

Although our study highlights how different types of cortex can be understood through the emergence of functional differentiation across space, it also raises a number of key questions for future research into how topography shapes function. First, although our study shows that association and unimodal cortex systematically vary in how function changes across the surface of the brain, this metric does not discriminate between systems that are known to be distinctive in their functions. For example, although the variograms for both the fronto- parietal and default mode networks are similar, the situations in which these systems contribute to cognition are different. Likewise, the variograms in motor and visual cortex are similar, yet these systems have clear functional differences. It is likely that the different roles that these systems play in cognition may arise, not from the general way that function changes with space in these areas of cortex, but in terms of the specific location that these systems inhabit within the broader cortical landscape. In this way our study highlights the more abstract properties that distinguish association and unimodal cortex, but do not provide a concrete explanation for how these systems contribute to cognition and behaviour in a distinctive manner. Second, our study does not constrain accounts of why association and unimodal cortex have differences in the spatial differentiation that we observe. Our analysis highlights that microstructural differences, via a proxy of intracortical myelination, systematically track differences in the empirical variograms. However, there are likely to be multiple microstructural features that track these differences, and these microstructural properties may also vary as a consequence of experience. Therefore, it is important for future work to examine the different genetic and experiential changes that influence how function varies as a function of distance in both primary and association cortex to fully understand the influences that determine this fundamental feature of cortical organisation.

## Methods

### Imaging Data

The majority of the analyses were performed on the first 51 participants’ resting state fMRI from the Human Connectome Project’s minimally pre-processed dataset; this involved registration to a common MNI152 template, minimal spatial smoothing and extensive filtering for slow drifts, motion and other nuisance signals estimated using independent components analysis (28). The 4D fMRI datasets for each participant were projected onto the Conte32k surface and the number of faces reduced resulting in 10,000 remaining vertices (using Matlab’s reducepatch command). Two resting-state runs (with opposite phase encoding direction, left-to-right and right-to-left) were taken from each participant. No further pre-processing was performed on the data.

Group averaged data from 14 macaque monkeys was used from the Newcastle cohort. Surface geodesic distance and homologous regions to the human data were taken from (29).

The vertex-wise map of cortical myelin was the group-average map taken from the Human Connectome Project 900-subject release; it is released in the Conte32k surface space and reduced to the same 10,000 vertices as the fMRI data. Similarly, the Yeo cortical parcellation (4) in Conte32k surface space was taken from the same HCP 900 data release and was also reduced to 10,000 vertices. The 50 Neurosynth data derived topic maps were downloaded in MNI152 2mm space and then projected onto the mid-thickness Conte32k surface using the Connectome Workbench (30) and then reduced to the same 10,000 vertices. Topics that were not related to cognitive tasks/states were removed, leaving 24 topics.

### Geodesic distance

Pairwise geodesic distance was calculated along the cortical surface between all vertices (excluding the medial wall) using the Connectome Workbench tools, as implemented through the BrainSmash toolbox (31). This was done on each hemisphere’s mid-thickness Conte32k surface reduced to 10,000 vertices prior to calculating the distances. The resulting vertex-wise distance matrices were used in all subsequent analyses.

### Functional connectivity

The functional connectivity affinity matrix was first calculated between all 10,000 vertices for each individual fMRI scan using Pearson’s correlation between the BOLD time series. For group-average results, the correlation coefficients were subsequently Fisher transformed and then for each vertex, averaged across subjects before applying an inverse Fisher transform, resulting in values between -1 and 1 for each edge of the functional connectivity matrix. Using a bounded similarity metric (0 = no similarity, 1/-1 identical) aids comparison across individuals/vertices and facilitates interpretation for the resulting empirical variograms.

### Empirical variograms

The empirical variogram was calculated by quantifying how functional connectivity decreases in similarity as distance increases. To do this, all distances between pairs of vertices were collapsed into 20 equally spaced bins. Subsequently, the difference in functional connectivity (Pearson’s correlation coefficient) between pairs of vertices was calculated and formed into equally spaced bins using a Gaussian smoothing function (following the approach set out in (31, 32)). This resulted in a whole-cortex empirical variogram. For vertex-wise variograms, the same approach was taken but repeated for every row of the functional connectivity/distance matrix separately, resulting in a simplified form of the empirical variogram for each vertex.

### Theoretical variogram

It is common practice to fit a function to empirical variograms, this is typically used prior to spatial regression; however, in our case, it allows us to compactly summarise the shape of the empirical variogram with a small number of parameters, facilitating comparisons across datasets and vertices, and aggregation across multiple vertices. For the reported analyses we used an exponential function. This is motivated by a range of prior studies suggesting exponential relationships between distance and various neural measures (e.g., (33)). We also performed a similar fit for two other theoretical models (a Gaussian and a periodic model which allows for non-monotonic functions), with qualitatively similar results. Empirical variograms were trimmed to bins between 2 and 19 (to remove bins with few sampled distances). Subsequently, non-linear least squares was used to estimate sill and the range.

### Low-dimensional embedding of functional connectivity

The principal connectivity gradient was calculated using the Brainspace toolbox (34). This involved taking the group-average functional connectivity affinity matrix and performing non-linear dimensionality reduction using the Laplacian Eigenmaps approach, separately for each hemisphere.

### Clustering

Agglomerative hierarchical clustering, with ward linkage and the Euclidean distance metric was applied simultaneously to all the vertex-wise variograms separately for each cortical hemisphere. Subsequently, SciPy’s *fcluster* command was used to flatten the hierarchy into two clusters. To assess the robustness of the resulting clusters each vertex’s variogram was correlated with all other variograms calculated in a separate fMRI run within the same individual. The correlation scores were Fisher transformed and then subsequently averaged both within and across clusters.

### Cognitive tasks

From the Neurosynth 50 data-derived topics dataset (35), those that did not refer to cognitive or behavioral states were removed, leaving: cognitive, inhibition, motor, numerical, action, conflict, spatial, emotion, empathy, decision, pain, memory, language, semantic, face, imagery, visual, eye movement, motion, attention, auditory, reward, social, working memory. The corresponding map for each topic was thresholded (absolute value z>10) and binarized, resulting in a vertex-wise mask of values that were strongly implicated for that topic (other thresholds produced qualitatively similar results). For each topic, the range and sill (taken from the theoretical variogram from the group average functional connectivity analysis) for each vertex within each mask were averaged together.

### Myelin

The estimated intracortical myelin maps derived from the ratio of T1 and T2 weighted MR images (16) from the Human Connectome Project were split into deciles based on their estimated myelin level. The empirical variograms of vertices within each decile were averaged. In addition, the estimated average myelin value for each of the clusters (see above) were calculated.

Python code to reproduce the analyses is available <u>here</u>

## Notes

### Competing Interest Statement

The authors have declared no competing interest.

## References

1. M. M. Mesulam, From sensation to cognition. Brain 121, 1013–1052 (1998).

2. D. J. Felleman, D. C. Van Essen, Distributed hierarchical processing in the primate cerebral cortex. Cereb Cortex 1, 1–47 (1991).

3. D. S. Margulies, et al., Situating the default-mode network along a principal gradient of macroscale cortical organization. Proceedings of the National Academy of Sciences 113, 12574–12579 (2016).

4. B. T. Thomas Yeo, et al., The organization of the human cerebral cortex estimated by intrinsic functional connectivity. J Neurophysiol 106, 1125–1165 (2011).

5. R. M. Braga, D. J. Sharp, C. Leeson, R. J. S. Wise, R. Leech, Echoes of the brain within default mode, association, and heteromodal cortices. Journal of Neuroscience 33, 14031–14039 (2013).

6. R. M. Braga, R. L. Buckner, Parallel Interdigitated Distributed Networks within the Individual Estimated by Intrinsic Functional Connectivity. Neuron 95, 457-471.e5 (2017).

7. R. B. H. Tootell, M. S. Silverman, E. Switkes, R. L. De Valois, Deoxyglucose Analysis of Retinotopic Organization in Primate Striate Cortex. Science 218, 902–904 (1982).

8. M. I. Sereno, et al., Borders of Multiple Visual Areas in Humans Revealed by Functional Magnetic Resonance Imaging. Science 268, 889–893 (1995).

9. M. I. Sereno, R.-S. Huang, A human parietal face area contains aligned head-centered visual and tactile maps. Nat Neurosci 9, 1337–1343 (2006).

10. E. Formisano, et al., Mirror-Symmetric Tonotopic Maps in Human Primary Auditory Cortex. Neuron 40, 859–869 (2003).

11. M. E. Raichle, The Brain’s Default Mode Network. Annu. Rev. Neurosci. 38, 433–447 (2015).

12. J. Smallwood, et al., The default mode network in cognition: a topographical perspective. Nature Reviews Neuroscience 22, 503–513 (2021).

13. S. Oligschläger, et al., Gradients of connectivity distance are anchored in primary cortex. Brain Struct Funct 222, 2173–2182 (2017).

14. Statistics for Spatial Data - Noel Cressie - Google Books (September 2, 2022).

15. D. A. Leopold, Y. Murayama, N. K. Logothetis, Very Slow Activity Fluctuations in Monkey Visual Cortex: Implications for Functional Brain Imaging. Cerebral Cortex 13, 422–433 (2003).

16. M. F. Glasser, D. C. V. Essen, Mapping Human Cortical Areas In Vivo Based on Myelin Content as Revealed by T1- and T2-Weighted MRI. J. Neurosci. 31, 11597–11616 (2011).

17. S. R. Quartz, The constructivist brain. Trends in Cognitive Sciences 3, 48–57 (1999).

18. J. Freeman, E. P. Simoncelli, Metamers of the ventral stream. Nat Neurosci 14, 1195–1201 (2011).

19. J. R. Stroop, Studies of interference in serial verbal reactions. Journal of Experimental Psychology 18, 643–662 (1935).

20. A. Baddeley, Working Memory. Science 255, 556–559 (1992).

21. J. Duncan, The multiple-demand (MD) system of the primate brain: mental programs for intelligent behaviour. Trends in Cognitive Sciences 14, 172–179 (2010).

22. D. L. Schacter, D. R. Addis, The cognitive neuroscience of constructive memory: remembering the past and imagining the future. Philosophical Transactions of the Royal Society B: Biological Sciences 362, 773–786 (2007).

23. M. A. L. Ralph, E. Jefferies, K. Patterson, T. T. Rogers, The neural and computational bases of semantic cognition. Nat Rev Neurosci 18, 42–55 (2017).

24. ,R. L. Buckner, L. M. DiNicola, The brain’s default network: updated anatomy, physiology and evolving insights. Nat Rev Neurosci 20, 593–608 (2019).

25. P. S. Goldman-Rakic, M. L. Schwartz, Interdigitation of contralateral and ipsilateral columnar projections to frontal association cortex in primates. Science 216, 755–757 (1982).

26. J. Duncan, An adaptive coding model of neural function in prefrontal cortex. Nat Rev Neurosci 2, 820–829 (2001).

27. C. J. Bajada, J. Schreiber, S. Caspers, Fiber length profiling: A novel approach to structural brain organization. NeuroImage 186, 164–173 (2019).

28. G. Salimi-Khorshidi, et al., Automatic denoising of functional MRI data: Combining independent component analysis and hierarchical fusion of classifiers. NeuroImage 90, 449–468 (2014).

29. T. Xu, et al., Interindividual Variability of Functional Connectivity in Awake and Anesthetized Rhesus Macaque Monkeys. BPS: CNNI 4, 543–553 (2019).

30. D. S. Marcus, et al., Human Connectome Project informatics: Quality control, database services, and data visualization. NeuroImage 80, 202–219 (2013).

31. J. B. Burt, M. Helmer, M. Shinn, A. Anticevic, J. D. Murray, Generative modeling of brain maps with spatial autocorrelation. NeuroImage 220, 117038 (2020).

32. J. Viladomat, R. Mazumder, A. McInturff, D. J. McCauley, T. Hastie, Assessing the significance of global and local correlations under spatial autocorrelation: A nonparametric approach. Biometrics 70, 409–418 (2014).

33. M. Shinn, et al., Spatial and temporal autocorrelation weave complexity in brain networks. 2021.06.01.446561 (2022).

34. R. Vos de Wael, et al., BrainSpace: a toolbox for the analysis of macroscale gradients in neuroimaging and connectomics datasets. Communications biology 3, 1–10 (2020).

35. T. Yarkoni, R. A. Poldrack, T. E. Nichols, D. C. Van Essen, T. D. Wager, Large-scale automated synthesis of human functional neuroimaging data. Nat Methods 8, 665–670 (2011).

